# Is most published research really false?

**DOI:** 10.1101/050575

**Authors:** Jeffrey T. Leek, Leah R. Jager

## Abstract

There has been an increasing concern that most published medical findings are false. But what does it mean to be false? Here we describe the range of definitions of false discoveries in the scientific literature. We summarize the philosophical, statistical, and experimental evidence for each type of false discovery. We discuss common underpinning problems with the scientific and data analytic practices and point to tools and behaviors that can be implemented to reduce the problems with published scientific results.

## 1 Introduction

Most published research findings are false.

This statement seems absurd on the first reading. Scientific research is carried out by highly trained and skilled scientists, vetted through peer review, and publicly scrutinized once it appears in journals. The entire scientific publishing infrastructure was originally conceived to prevent the publication of incorrect results and provide a forum for correcting false discoveries [Csiszar, 2016]. It seems inconceivable that most of the findings that pass through this process are false.

But this system was invented before modern computing, data generation, scientific software, email, the internet, and social media. Each of these inventions has placed strain on the scientific publication infrastructure. These modern developments have happened during the careers of practicing scientists. Many laboratory leaders received their training before the explosion of cheap data generation, before the widespread use of statistics and computing, and before there was modern data analytic infrastructure [Irizarry, 2012]. At the same time, there has been increasing pressure from review panels, hiring committees, and funding agencies to publish positive and surprising results in the scientific literature. These trends have left scientists with a nagging suspicion that some fraction of published results are at minimum exaggerated and at worst outright false.

This suspicion was codified in a widely read paper titled, “Why most published research findings are false” [Ioannidis, 2005b]. The paper focuses on how application of the hypothesis testing framework could result in a preponderance of false positive results in the medical literature. The key argument is analogous to population based screening using a biomarker. A very sensitive and specific biomarker may still have a low positive predictive value if the disease is not prevalent in the population. Analogously suppose that the null hypothesis is true for 99% of the hypotheses being considered by scientific investigators. If scientists test 1,000 hypotheses with a statistical power of 80% then 1, 000 × 1% × 80% = 8 true alternative hypotheses should be correctly detected. Even though the Type I error rate is much lower, the prevalence of null hypotheses is much higher. With a Type I error rate of 5% then we expect 1, 000×99%×5% = 49.5 null hypotheses will be incorrectly detected. In this situation 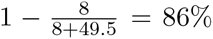 of rejected hypotheses will actually be null. Assuming selective reporting of positive results, a lower prevalence of true alternatives in hot fields, or other sources of bias in the publication process makes this estimate even worse.

Under this argument a “false” research finding is defined as a finding where the null hypothesis is true, but incorrectly called statistically significant by a researcher. At its core is an argument about the way statistical evidence is assessed across the medical literature. But this is hardly the only difficulty with the modern scientific process. There are a variety of other ways that a statistical result can be considered false or suspect. It is possible to confuse correlation with causation; a predictive model may overfit the training data; a study may be underpowered; results may be over-interpreted or misinterpreted by the scientific press [Leek and Peng, 2015a, Leek and Peng, 2015b]. Each of these scenarios may undermine the credibility of a scientific result without satisfying the definition of false from a hypothesis testing perspective.

The added complexity of the scientific process also contributes to the difficulties in verifying the results from scientific publications. It used to be sufficient to describe scientific results using text and figures, but it is increasingly necessary to describe a set of scientific results through scientific code, by releasing a data set, or through a series of complex protocols that can not be completely described in text [Peng, 2011]. Since this change has happened during the careers of many active scientists, there has been a slow realization that publications frequently do not provide sufficient detail to describe the scientific and computational protocols included in a study.

This complexity raises the question of how often it is possible to reproduce or replicate the results of a particular scientific study. *Reproducibility* is defined as the ability to re-create all of the figures and numbers in a scientific publication from the code and data provided by the authors. *Replicability* is defined as the ability to re-perform the experiments and computational analyses in a scientific publication and arrive at consistent results. Both of these ideas are related to the rate of “false discoveries” in the medical literature. If a study can not be reproduced, then it is impossible to fully assess whether the evidence supports any claims of statistical significance. If a study does not replicate, then it raises the question of whether the original report was a false discovery.

This review tackles each of these challenges separately and attempts to summarize what we know about the current state of false discoveries in science. We also point to ongoing and potential initiatives to reduce the statistical problems within the medical and scientific literature.

## 2 Defining false discoveries in the medical literature

Here we consider a range of potential definitions of issues in the medical literature, starting with computational issues and concluding with false discoveries at the completion of an analysis. Any single scientific or medical study consists of a question of interest, an experimental design, an experiment, data sets, analysis code, and conclusions.

We will consider the following issues with the medical literature:

- **Reproducibility** - A study is reproducible if all of the code and data used to generate the numbers and figures in the paper are available and exactly produce the published results.
- **Replicability** - A study is replicable if an identical experiment can be performed like the first study and the statistical results are consistent.
- **False discovery** - A study is a false discovery if the result presented in the study produces the wrong answer to the question of interest.

Reproducibility is the easiest of these problems to both define and assess. Assessing reproducibility involves checking the published manuscript, looking for published data and code, then comparing the results of that data and code to the published results. If they are the same the study is reproducible, if they are not, then the study is not.

Replicability is a more challenging concept to both define and measure. A study replicates if the same experiment can be performed a second time with consistent results. If the data collected during the study are subject to sampling variability then even in the best case scenario the results of a replication will not be identical to the original study. However, we would expect that the results would be within the range of values predicted by the parameter estimates and variability estimates from the original study. The difficulties in assessing replicability are compounded by potential for publication bias, regression to the mean, fragility of scientific results to a particular context, and imperfect replication.

A false discovery is the most challenging of these three problems to assess. A false discovery means that the reported parameter or answer to a scientific question is not consistent with the underlying natural truth being studied. A false discovery is the most difficult to assess because we rarely know the true state of nature for any particular scientific study. Single replications are not sufficient to separate true discoveries from false discoveries since both the original study and the replication are subject to sampling error and other potential difficulties with replication studies. Repeated replications or near replications that all point to a similar conclusion are the best way to measure false discoveries in the medical literature. However, repeated replication or near replication of identical studies is very expensive and tends to only occur for highly controversial ideas - such as the claim that vaccines cause autism.

## 3 What is the rate of reproducibility of the scientific literature?

We begin by considering the rate of reproducibility of scientific studies. Computational reproducibility was first identified as a key component of the scientific process over three decades ago within the computational and statistical community [Buckheit and Donoho, 1995]. But the role of reproducibility in the broader scientific process was not highlighted until the mid-2000s across a variety of fields from Biostatistics [Peng, 2009] and Epidemiology [Peng et al., 2006] to Physics [Buckheit and Donoho, 1995]. Ultimately the reproducibility of scientific results became a baseline standard by which the data analysis in any particular study should be judged.

Prior to the widespread knowledge of the importance of reproducible research many papers did not provide data and code. Part of this issue was cultural - it was not well known that providing data and code was a key component of the scientific publication process. This issue was compounded since most scientists were not trained in statistics and computation. Moreover, the tools for creating and sharing reproducible documents were often difficult to use for people without sufficient computational training.

There have been multiple studies evaluating reproducible research across different disciplines.

- **Study** Repeatability of published microarray gene expression analyses [Ioannidis et al., 2009] **Main idea** This paper attempts to collect the data used in published papers and to repeat one randomly selected analysis from the paper. For many of the papers the data was either not available or available in a format that made it difficult/impossible to repeat the analysis performed in the original paper. The types of software used were also not clear. **Important drawback** This paper focused on 18 data sets in 2005-2006. This paper was published early in the era of reproducibility and so is potentially driven by cultural change in genomics studies 10 years ago.
- **Study** Toward reproducible computational research: An empirical analysis of data and code policy adopted by journals [Stodden et al., 2013] **Main idea** The authors evaluated code and data sharing policies for 170 journals. They found that 38% and 22% of these journals had data and code sharing policies, respectively. **Important drawback** The chosen journals were more computationally focused, although also included high-impact journals such as Nature Genetics, Cell, and Lancet.
- **Study** Believe it or not: how much can we rely on published data on potential drug targets? [Prinz et al., 2011] **Main idea** 67 studies that focus primarily on oncology produce relevant code and data only 20% of the time. **Important drawback** The data, code, and methods used to perform this study are not available and so it is not a scientific study.
- **Paper** Next-generation sequencing data interpretation: enhancing reproducibility and accessibility [Nekrutenko and Taylor, 2012] **Main idea** As part of an opinion piece, the authors randomly selected 50 of 378 papers in 2011 that used the same alignment technique. Of these papers, only 7 provided all necessary details for reproduciblity. Nineteen provided enough information on software implementation and almost half provided access to data. **Important drawback** The set of papers examined focuses on a relatively small area of science.
- **Study** Public availability of published research data in high-impact journals [Alsheikh-Ali et al., 2011] **Main idea** In a sample of 500 research articles from 50 high-impact journals in 2009, only 9% of articles made raw data fully available. Of the 50 journals surveyed, 44% had a policy explicitly requiring materials to be available as a condition of publication and 12% had no specific policy for data availability. Even when data-sharing policies were in place, more than half of the sampled articles did not meet the journal’s stated requirements for data availability. **Important drawback** Data availability is only one piece of reproducibility; to be fully reproducible code must be available as well.
- **Study** Case studies in reproducibility [Hothorn and Leisch, 2011] **Main idea** 100 randomly sampled papers from Bioinformatics were sampled and evaluated for code and data availability. Code for simulations was available more than 80% of the time, data was available around 50% of the time when used, and code was available slightly more than 60% of the time when used. **Important drawback** This paper represents a relatively solid evaluation of the rate of reproducibility, but only in the bioinformatics world where the rate of reproducibility might be artifically high.

In addition to these empirical evaluations, there has been a large literature dedicated to philosophical and opinion based pieces on reproducibility in the sciences. There is likely some truth to these opinion pieces, but they tell us little about the actual rate that studies are not reproducible. Certainly, the rate at which studies are reproducible varies by discipline. The most extensive studies of reproducibility have occurred within the Bioinformatics community, where the rate of computational and statistical sophistication is arguably higher than other disciplines. Research areas with a mature view of statistics and computation likely produce more reproducible research than areas newly introduced to computational and empirical research.

Lack of reproducibility itself does not certainly imply that a particular scientific result is false. A study may be fully correct including the reporting of a true positive and not be fully reproducible. On the other hand, a fully reproducible study may be riddled with errors and be wrong. Without full access to the data and code describing a particular scientific result it is difficult to assess how credible that result should be. Regardless, most effort to improve the scientific literature has so far focused on reproducibility since it is very difficult to evaluate replicability or true and false discoveries without data and code.

## 4 What is the rate of replication in the medical literature?

Reproducibility is concerned with obtaining the exact results of a published study once the code and data are available. A higher standard is the ability to replicate a published study. Replication involves following the published protocol and repeating the entire experiment, data collection, and data analysis process. The goal is to determine if the result of the replication is consistent with the original study.

Replication studies include several levels of subtlety. The first is defining what it means to be a successful replication. Definitions of successful replication have included consistent effect sizes, consistent distributions, and consistent measures of statistical significance. The difficulty is that both original and replication experiments involve multiple sources of variation. Sampling variation alone may explain why a result may be significant in one study and not another. In other cases, regression to the mean and publication bias may lead to reductions in the estimated effects in replication studies. Finally, replication studies are extremely hard to perform well since variations in measurement technology, population characteristics, study protocols, and data analyses may lead to non-replication.

Despite the expense of performing replication of scientific experiments, there is increasing energy and attention being focused on these studies. Typically only the most important and widely publicized studies have been the focus of replication, but that is changing with larger scale studies of replication and increased incentives to perform this type of research. Here we focus on a subset of important replication studies.

- **Study** Estimating the reproducibility of psychological science [Collaboration et al., 2015] **Main idea** The goal of this Reproducibility Project: Psychology study is to re-perform 100 experiments in psychology and evaluate the replicability of results in psychological science. The claim is that many of the results are not replicable; the study found 36% of the experiments were significantly replicated. **Important drawback** The study only performs a single replication of each study with potential issues due to power, bias, and study design.
- **Study** What should we expect when we replicate? A statistical view of replicability in psychological science [Leek et al., 2015] **Main idea** A re-analysis of the psychological science replication data shows that 77% of the replication effects fall within the 95% prediction interval for the effect size based on the original study. **Important drawback** This paper does not model the potential biases in either the original or replication studies. Consistency of effect estimates does not say anything about whether the results are true or false discoveries.
- **Study** Contradicted and initially stronger effects in highly cited research. [Ioannidis, 2005a] **Main idea** This paper looks at studies that attempted to answer the same scientific question where the second study had a larger sample size or more robust (e.g. randomized trial) study design. Some effects reported in the second study do not match the results exactly from the first. **Important drawback** The title does not match the results. 16% of studies were contradicted (meaning effect in a different direction). 16% reported smaller effect size, 44% were replicated and 24% were unchallenged. So 44% + 24% + 16% = 86% were not contradicted. Lack of replication is also not proof of error.
- **Study** A survey on data reproducibility in cancer research provides insights into our limited ability to translate findings from the laboratory to the clinic [Mobley et al., 2013] **Main idea** Researchers surveyed faculty/trainees at MD Anderson Cancer Center about their ability to replicate published results. Of those who responded, 55% reported that they had tried and failed to replicate a result from a published paper. For those who contacted the authors of the original paper, less than 40% received a positive/helpful response in reply. **Important drawback** Voluntary sample with a low response rate; participants all from a single institution.
- **Study** Investigating variation in replicability: The “Many Labs” replication project [Klein et al., 2014] **Main idea** The paper considered 13 published high-profile results and used multiple labs to replicate the results. They successfully replicated 10 out of 13 results and the distribution of results allows us to measure the potential effect of regression to the mean. **Important drawback** The paper covers only 13 high-profile experiments in psychology. This is by far the strongest, most comprehensive, and most reproducible analysis of replication among all the papers surveyed here.

These replication efforts suggest that some studies replicate and some studies do not. The best estimates of replication are available for the 13 studies in the “Many Labs” replication project - where we can estimate the impact of differing study designs, experimental procedures, and personnel. In that case 10/13 studies replicated, however this represents a relatively narrow area of psychology and only the most high profile results. The replication rate is again likely field specific. In psychology, where most of the replication efforts have been concentrated, the true rate of replication likely lies between the 36% of studies with statistically significant results in the same direction in the Reproducibility Project: Psychology and the 77% of studies that replicated in the “Many Labs project”.

The reality is that consistent replication of all scientific studies is not a viable approach. Currently, only papers that garner sufficient interest and attention in the form of high-profile publication or press will be subject to replication. An alternative strategy would be to perform random audits of specific disciplines or sub-disciplines, like the “Many Labs” project, to provide an overall estimate of the replication rate for a particular field.

## 5 What is the rate of false discoveries in the medical literature?

Replication represents an attempt to confirm knowledge that we think we have garnered from a scientific study. But a single replication of a scientific result is not sufficient to measure the true state of the world. If only a single replication is performed, there is variation in both the original study and in the replication study. So if the results are not consistent we can’t be sure if that is because either study has produced a false result or if it is simply expected due to natural variation.

The problem of the rate of false discoveries in the medical literature has primarily been addressed through theoretical arguments and discussion. Only recently have efforts been made to empirically estimate the rate of false discoveries in the scientific or medical literature. Some of the key studies in the area include the following.

- **Paper** Why most published research findings are false [Ioannidis, 2005b] **Main idea** People use hypothesis testing to determine if specific scientific discoveries are significant. This significance calculation is used as a screening mechanism in the scientific literature. Under assumptions about the way people perform these tests and report them it is possible to construct a universe where most published findings are false positive results. **Important drawback** The paper contains no real data, it is purely based on a theoretical argument about the behavior of scientists and how they choose hypotheses to study.
- **Study** Drug development: Raise standards for preclinical research [Begley and Ellis, 2012] **Main idea** Many drugs fail when they move through the development process. Amgen scientists tried to replicate 53 high-profile basic research findings in cancer and could only replicate 6. **Important drawback** This is not a scientific paper. The study design, replication attempts, selected studies, and the statistical methods to define “replicate” are not defined. No data is available or provided.
- **Study** An estimate of the science-wise false discovery rate and application to the top medical literature. [Jager and Leek, 2014] **Main idea** The paper collects *p*-values from published abstracts of papers in the medical literature and uses a statistical method to estimate the false discovery rate proposed in [Ioannidis, 2005b] above. This paper estimates the rate of false discoveries at 14%. **Important drawback** The paper only collected data from major medical journals and the abstracts; *p*-values can be manipulated in many ways that could call into question the statistical results in the paper.
- **Study** Investigating variation in replicability: The “Many Labs” replication project [Klein et al., 2014]. **Main idea** The paper considered 13 published high-profile results and used multiple labs to replicate the results. They successfully replicated 10 out of 13 results and the distribution of results allows us to measure the potential effect of regression to the mean. **Important drawback** The paper covers only 13 high-profile experiments in psychology. This is by far the strongest, most comprehensive, and most reproducible analysis of replication among all the papers surveyed here.

Only the study of published *p*-values [Jager and Leek, 2014] and the many labs replication project [Klein et al., 2014] empirically evaluate the rate of false discoveries in the medical literature. The study of published *p*-values suffers from potential issues with publication and other biases [Gelman and O’Rourke, 2014] - but is one of the only comprehensive estimates of the science-wise false discovery rate that is available. This study suggested that the rate of false discoveries in the medical literature was inflated, but only to 14%.

The “Many Labs” project represents a much higher standard for evaluating the rate of false discoveries in the literature. With repeated replications by multiple groups, it is possible to distinguish effects that are consistent from those that are not. This study suggested 24% of results were false discoveries. However, this approach to evaluating the veracity of the literature is time consuming and expensive, and we only have this estimate for a small sample of psychology experiments.

Overall the empirical rate of false discoveries in the medical literature is unknown. Our only empirical estimates - however shaky or specific they may be - suggest that most published research is not false. A key challenge for the future is designing studies that are rigorous, cover a broad fraction of important scientific disciplines, and are inexpensive and efficient enough to be performed regularly.

## 6 What is the rate of incorrectly analyzed data in the medical literature?

Among all studies that are replicable and reproducible, the primary source of false discoveries is likely due to incorrect experimental design or data analysis [Leek and Peng, 2015a]. There have been two very public examples of incorrect analysis producing questionable or false results. The first involves a series of papers that claimed to identify genomic signatures that could predict the response of patients to chemotherapy [Potti et al., 2006]. These studies were originally not reproducible. The irreproducibility of these studies has been a major source of discussion [Baggerly and Coombes, 2009, Baggerly, 2010, Micheel et al., 2012]. Reproducible versions of the analyses in the studies revealed major flaws in the statistical methods and computational software used to make the predictions [Baggerly and Coombes, 2009]. These flaws included problems with study designs, incorrect probabilistic statements, and predictions subject to random error.

A second public example, in economics, involves a paper that suggested that a high debt to GDP ratio was related to decreased economic growth [Rogoff and Reinhart, 2010]. The paper was ultimately used by politicians and economists as justification for austerity measures around the globe. A graduate student discovered an error in the spreadsheet used by the authors and this error received both widespread academic and press attention. However, the major issue with the analysis was the extremely limited sample size, choices about which data points to exclude, and unconventional weighting schemes.

These cases highlight the issues that arise with data analysis in nearly every paper in the scientific literature. As data become increasingly common and complicated, so too do the software and statistical methods used to analyze the data. There is increasing concern about the use of *p*-values, under powered studies, potential confounders, the winner’s curse, robustness of analytic pipelines, researcher degrees of freedom, and *p*-value hacking in scientific studies. There have now been studies focused on evaluating the extent of these issues in scientific and medical studies [Aschwanden and King, 2015].

There is a clear connection between poorly designed or analyzed studies and both a lack of replication and an increased risk of false discoveries. But we know little about the data analytic practices used across individual studies. There is some indication that certain fields tend to have low-powered and observational studies [Button et al., 2013]. Other fields have focused on the use of well-designed randomized trials. In either case, as data become more complicated, the role of data analytic pipelines on the ultimate validity of statistical results is not well understood. Understanding and improving the data analytic process still remains the surest way to reduce false discoveries in the medical literature.

## 7 How can we improve the scientific and medical literature?

Science is most likely not in the midst of a crisis of reproducibility, replicability, and false discoveries. But there are a large number of scientific studies that do suffer from the underlying computational and statistical issues [Aschwanden and King, 2015]. Addressing the real statistical problems in science requires efforts to improve the incentive system for scientists, to produce tools that simplify computation and statistics, and to increase training in computation, software development, and data analysis [Ioannidis, 2014].

### 7.1 Improving incentives

One of the major barriers to reproducibility, replicability, and true discoveries in science is an imbalanced incentive system. There is strong pressure to publish only “positive results” and to not publish replication studies. There is pressure against re-using published data, with strong advantages for data generators over data analysts. To improve the statistical side of science we need incentive systems to be aligned with solid statistical efforts.

Some incentive programs that are underway are the following.

- **The pre-registration challenge** [Cohoon, 2015] - provides researchers with an incentive of $1,000 to pre-register their hypotheses before performing a study. These studies will avoid p-hacking, outcome switching, and other potential problems with discovery based research.
- **Transparency and openness promotion guidelines** [Nosek et al., 2015] - these guidelines, adopted by hundreds of journals, commit these journals to promoting standards of openness, data sharing, and data citation.
- **NSF and NIH research output definitions** [Collins and Tabak, 2014, NIH, 2015, NSF, 2015] - both the NSF and NIH have recently made efforts to legitimize research outputs beyond academic publications including software production, data generation, and scientific outreach activities.

These initiatives represent excellent efforts to improve the incentive system for scientists who wish to perform reproducible, replicable, and accurate research. However, the major remaining impediment to the incentive system is the value placed on specific research activities by hiring, promotion, and grant-evaluation committees. These committees hold incredible sway over the careers of all scientists, but particularly the junior scientists who are most likely to adopt practices that avoid the issues we have discussed here.

The highest impact potential incentive would be for these committees to publicly and formally recognize the value in activities that take away time from publication but contribute to good science [Allen and Leek, 2013]. Specifically, by placing emphasis on behaviors like software maintainence, data generation and sharing, and peer review, these committees could have an immediate and dramatic impact on the rate of reproducibility, replicability, and accuracy in the scientific literature.

### 7.2 Improving tools

One of the key impediments to performing reproducible and replicable research in the past was the lack of availability of tools that make it easy to share data, to share code, and to perform correct analyses. Fortunately there has been an explosive growth in the availability of tools for these purposes over the last ten years. For general purpose analysis, the growth of the R programming language and the Python for Data Analysis community has lead to a host of free software for performing a wide range of analyses.

There are now a number of tools that facilitate distributing reproducible analysis code and pipelines.

- **knitr/rmarkdown** [Xie, 2015, Allaire et al., 2015] - these are R packages that have been developed for creating documents that interweave text, code, and figures. These documents are fully reproducible within the R environment and can be easily created by students with only basic R training.
- **iPython notebooks** [P´erez and Granger, 2007] - these create interactive and static documents that interweave text, code, and figures. These documents are fully reproducible and are supported by Github (a popular code sharing repository) so that they can be easily viewed online.
- **Galaxy** [Goecks et al., 2010] - this is an infrastructure for creating reproducible work flows by combining known tools. Galaxy workflows are reproducible and can be shared with other researchers.

There are similarities and differences across these different platforms, but the commonality is that they add negligible effort to an analysts data analytic workflow. Both knitr and iPython notebooks have primarily increased reproducibility among scientists who have some scripting experience. A major reason for their popularity is that code is written as usual but simply embedded in a simple-to-use document. The workflow isn’t changed for the analysts because they were going to write the code anyway. The platform just allows it to be included into an easily shareable document.

Galaxy has increased reproducibility for a range of scientists, but the primary user is someone who has less scripting experience. The Galaxy project has worked hard to make it possible for everyone to analyze data reproducibly. The reproducibility is almost incidental - a user would have to stitch pipelines together anyway, and Galaxy provides an easy way to do so. Reproducibility comes as an added bonus.

Data can now also be easily shared. Researchers used to post data to their personal websites, but it was largely impermanent in that state. When researchers changed institutions or web servers, data often was lost. Link rot has been one of the most fundamental sources for lack of reproducibility in science. Now, however, there are a variety of permanent places data can be made publicly available. General purpose sharing sites include the following.

- **Figshare** [Hannel, 2015] - if designated as publically available, an unlimited amount of data can be posted free of charge. Private data storage requires a fee. Figshare accepts all data types and provides a digital object identifier (doi) so data so can be cited.
- **Open Science Framework** [OSF, 2015] - can be used to post all types of data sets and can be integrated with Figshare.
- **Dataverse** [King, 2007] - free data storage hosted by Harvard.

Data can also be hosted in a variety of field-specific data repositories. Specific repositories have also been developed for handling sensitive data, such as personally identifiable information. As a result, there are resources available for all but the largest data-sharing efforts.

### 7.3 Improving data analysis

Even with the improvements in incentives and tools, a major obstacle to improving the issues with statistics in science involves providing sufficient training to the students and postdocs who perform most data analyses. It is no longer feasible to expect that all data analysis will be performed by a person with advanced training in statistics and data analysis. Every lab across every scientific discipline is now engaged in the generation of abundant and cheap data. There are simply not enough data analysts with advanced training to keep pace.

Several initiatives have risen to meet this demand. Some are short, intensive workshops in software engineering and data analysis. Others are online courses that can be used to dramatically scale basic training in these areas.

- **Software carpentry** [Wilson, 2006] - workshops on reproducible research, software design, and software use that are run at locations around the world.
- **Data carpentry** [Teal et al., 2015] - workshops on data cleaning, management, and analysis that are run at locations around the world.
- **Massive open online courses** (MOOCs) [Gooding et al., 2013] - A range of data science, data analysis, and reproducible research MOOCs that are available to all researchers around the world.

Materials and courses are now available to learn these critical techniques and improve the scientific process. Unfortunately, however, many scientific programs, including all medical education programs, do not require a single statistics or data analysis class. Given how frequently data and statistical analysis play a role in research that is published in the medical literature, the key next step is to integrate these training modules into the formal education of all scientists.

## 8 Conclusions

We have summarized the empirical evidence that most published research is not reproducible, replicable, or false. Our summary suggests that the most extreme opinions about these issues likely overstate the rate of problems in the scientific literature. However, there is clearly work to be done. Statistics and data analysis now play a central role across all sciences, including medical science. We need to work to encourage the adoption of best practices and available tools to improve the accuracy of published scientific results.

## 9 Acknowledgements

Parts of the text of this review have appeared in the blog posts: “A summary of the evidence that most published research is false” [Leek, 2013] and “Why the three biggest positive contributions to reproducible research are the iPython Notebook, knitr, and Galaxy” [Leek, 2014] and the book “How to be a modern scientist” [Leek, 2016].

